# Assessments of hepatitis B virus-like particles and Crm197 as carrier proteins in melioidosis glycoconjugate vaccines

**DOI:** 10.1101/2020.04.08.031658

**Authors:** Marc Bayliss, Matthew I. Donaldson, Giulia Pergolizzi, Andrew E. Scott, Sergey A. Nepogodiev, Lucy Beales, Michael Whelan, William Rosenberg, Hadrien Peyret, George P. Lomonossoff, Nicholas J. Harmer, Tim Atkins, Robert A. Field, Joann L. Prior

## Abstract

The Tier 1 select agent *Burkholderia pseudomallei* is the causative agent of melioidosis, a global pathogen and a major cause of pneumonia and sepsis for which no licensed vaccines currently exist. Previous work has shown the potential for *Burkholderia* capsular polysaccharide (CPS) to be used as a vaccine antigen but the T-cell independent nature of the immune response to this molecule requires the use of this polysaccharide as a glycoconjugate for vaccination. Recent studies have focussed on the use of Crm197 (a non-toxic mutant protein derived from diphtheria toxin) as the carrier but there are concerns regarding its potential to cause interference with other vaccines containing Crm197. Therefore research with alternative carrier proteins would be beneficial. In this study, CPS was isolated from the non-pathogenic *B. thailandensis* strain E555. This was chemically conjugated to Crm197, or Tandem Core™ virus-like particles (TCVLP) consisting of hepatitis B core protein, which is the first documented use of VLPs in melioidosis vaccine development. Analysis of CPS-specific IgG antibody titres showed that mice vaccinated with the Crm197 conjugate generated significantly higher titres than the mice that received TCVLP-CPS but both conjugate vaccines were able to protect mice against intraperitoneal *B. pseudomallei* strain K96243 challenges of multiple median lethal doses.

## Introduction

*B. pseudomallei* is the causative agent of melioidosis, a potentially lethal human and animal disease disseminated through soil and water [1, 2, 3]. *B. pseudomallei* is classified as a Tier 1 bio-threat agent by the US Centers for Disease Control and Prevention (CDC) [4] and it is estimated that the annual number of deaths resulting from melioidosis (89,000) is comparable to that of measles [5]. For these reasons, development of a *B. pseudomallei* vaccine is a priority. *B. pseudomallei* capsular polysaccharide (CPS), - 3-)-2-*O*-acetyl-6-deoxy-β-D-*manno*-heptopyranose-(−1 polymer, is a protective antigen and virulence determinant [6, 7]. The immune response to polysaccharides, which are generally T cell-independent type 2 antigens, can be significantly improved by conjugation to a carrier protein, which leads to the formation of carbohydrate-specific CD4+ T-cells which provide help to antibody producing B cells [8, 9]. It has been shown that mice vaccinated with conjugates utilising bovine serum albumin as a carrier for CPS or TetHc as a carrier for synthetically-synthesised CPS are significantly protected against non-inhalational *B. pseudomallei* challenge compared to controls, but sterilising immunity was not achieved in every animal [10, 11]. Recently, one of the most commonly used carrier proteins in licensed conjugate vaccines, Crm197 [12, 13], has been shown to protect mice against an inhalational challenge of *B. pseudomallei* and achieve high levels of sterilising immunity when conjugated to CPS [14]. This result and the obvious cost and safety advantages of Crm197 justify its use in melioidosis vaccine research but alternative carrier proteins should be sought due to concerns that prior exposure to a carrier can reduce carbohydrate-specific immune responses in other same-carrier-based conjugate vaccines. [9, 15, 16, 17, 18].

Alternative potential carrier proteins include virus-like particles which are formed from viral structural proteins, typically capsid or envelopes, which have the property of self-assembly for the formation of structures that mimic intact virus particles [19]. VLPs are non-infectious, non-replicating, and their particulate nature leads to efficient uptake by dendritic cells [20]. Antigenic epitopes in a VLP construct are displayed in a highly repetitive manner. This leads to B-cell receptor cross-linking and CD4+ and CD8+ T-cell stimulation, inducing both humoral and cellular immune responses [19, 21, 22], which are likely to be required for immunity to melioidosis [23]. Currently, recombinant VLP-based vaccines against Hepatitis B Virus (HBV), Human Papilloma Virus (HPV) and Hepatitis E Virus (HEV) have been approved and licensed for human use; numerous other VLPs designed to generate protection against other viral diseases are under study and/or clinical trial [24].

Hepatitis B core antigen (HBcAg) is an effective activator of macrophages, can act as both a T-cell dependent and T-cell independent antigen [25, 26] and readily assembles into VLPs that have been explored for therapeutic use [27]. Furthermore, they are attractive carrier proteins as foreign constructs can be inserted into the HBcAg protein, which result in strong immune responses to both VLP and insert [28, 29]. In order to facilitate the conjugation of CPS to the major immunodominant region (MIR) of the VLP surface, Tandem Core™ technology was introduced [30] (generating TCVLPs) (Figure S1). A Tandem Core™ is two HBcAg sequences genetically linked, which allows for insertion of a wider range of constructs whilst remaining assembly competent (Figure S2). In the current study, we have genetically inserted six lysine residues flanked on either side by three aspartic acid residues into MIRs of TCVLPs, which can be used for chemical conjugation to polysaccharide antigens, such as *Burkholderia* CPS (Figure S3).

Further, CPS conjugates of TCVLP and Crm197 were prepared and evaluated for immunogenicity and protective efficacy in a murine model of melioidosis. We show that both TCVLPs and Crm197 can be used as effective carrier proteins in CPS glycoconjugate vaccines for melioidosis despite the significant difference in CPS antibody titres generated between them.

## Materials and methods

### Bacteria/CPS isolation

The O-PS deficient mutant of *B. thailandensis* E555 harbouring a kanamycin-resistance marked, in-frame deletion of its *wbiI* gene (*B. thailandensis* E555 :: *wbiI* (p-Knock KmR)) [31] was grown in 2 L of LB broth overnight at 37°C with shaking. The CPS was extracted *via* a modified hot-phenol method and purified as described previously [32].

For animal challenges, *B. pseudomallei* K96243 was inoculated from a glycerol stock into 100 mL L-broth and incubated for 24 h at 37°C with shaking. The optical density (OD _590 nm_) was adjusted to 0.4, corresponding to approximately 4 × 10^8^ CFU/mL, and diluted in L-broth to the correct concentration for challenge.

### Production of the GD3K6D3G pEAQ-HT t-HBcAg plasmid

Building on work by Jegerlehner *et al.*, [33], who conjugated antigens to VLP to a single lysine inserted into the MIR sequence, a 14 amino acid peptide insert into the pEAQ-*HT t-HBcAg* plasmid was designed (Figure S3) that comprised Glycine-Aspartate3-Lysine6-Aspartate3-Glycine (GD3K6D3G). The oligo lysine sequence was to create multiple conjugation points for CPS, the aspartate residue flanks were to negate a potential positive charge imbalance in the E1-loop, and glycine spacers used to separate the insert sequence from the native peptide. Tandem Core™ technology was used since the introduction of the GD3K6D3G sequence into monomeric cores abrogated particle assembly [34]. To create the GD3K6D3G pEAQ-*HT t-HBcAg* plasmid, restriction digests (*Ase*I & *Sal*I, New England Biolabs) of the pEAQ-*HT t-HBcAg* plasmid [30] was carried out using standard conditions and the sample was run on a 0.8 % agarose gel. The corresponding band was excised from the gel and the plasmid extracted using Qiaquick gel extraction kit (Qiagen). The forward and reverse phosphorylated primers (Sigma Chemical, FWD: 5’-TCGACGGAGACGATGACAAGAAGAAGAAGAAGAAGGATGACGATGGTAT; REV:GCCTCTGCTACTGTTCTTCTTCTTCTTCTTCCTACTGCTACCATAAT-5’) were annealed and the ligation reaction was carried out using a 3:1 ratio of insert to plasmid backbone. After overnight incubation at 16 °C, Top10 chemically competent *E. coli* cells (Invitrogen) were transformed with the plasmid. Colony PCR and sequencing was used to confirm the successful cloning of the t-HBcAg GD3K6D3G plasmid.

### Expression of Tandem Core™ constructs in Nicotiana benthamiana

Heterotandem core (GD3K6D3G construct: MIR 1 empty and MIR 2 containing 6 x lysines flanked on each side by 3 x aspartic acid residues) were transformed into *Agrobacterium tumefaciens* LBA4404 by electroporation and propagated at 28°C in LB media containing 50 µg/mL kanamycin and 50 µg/mL rifampicin. Transient expression was carried out by agroinfiltration of 3 - 4 week old *Nicotiana benthamiana* leaves. *Agrobacterium tumefaciens* strains were sub-cultured and grown overnight, pelleted and resuspended to OD_600 nm_ = 0.4 in MMA (10 mM MES-NaOH, pH 5.6; 10 mM MgCl_2_; 100 mM acetosyringone) and then infiltrated into leaf intercellular spaces using a blunt-ended syringe. Plants were grown in a greenhouse maintained at 23 - 25°C and infiltrated 3 - 4 weeks after the seedlings were pricked out. The first four mature leaves of each plant were selected for infiltration. Plant tissue was harvested 6 days post infiltration.

The fresh plant material was weighed (100 g leaves harvested from 60 plants) and added to phosphate buffer [100 mM sodium phosphate; Roche complete protease inhibitor tablet (EDTA-free) as per manufacturer’s instructions] (3 mL per gram of plant material) and homogenised in a blender. Large debris was removed by centrifugation at 15,000 × g for 14 minutes and the supernatant filtered through a 0.45 µm syringe filter. The volume of the clarified lysate was then reduced from 380 mL to 180 mL on a rotavapor at 15°C. The supernatant was purified using a two-step sucrose cushion with 75 % (w/v) and 25 % (w/v) sucrose layers in ultracentrifuge tubes. The gradients were centrifuged at 240,000 × g for 2.5 h at 4°C. The sucrose layers were collected and dialyzed against PBS and analyzed by western blot. The VLP containing fractions were combined and extensively dialyzed (5 × 1 L) against ammonium bicarbonate (20 mM, pH 7.4) [35]. The fractions from the 75 % (w/v) and 75-25 % (w/v) interface of the sucrose gradient, containing most of the VLPs, were subjected to further purification on a Sephacryl S500 column over five runs. The first chromatography run was eluted into PBS and all subsequent runs were eluted into 20 mM ammonium bicarbonate buffer pH 7.4. Samples were analysed using transmission electron microscopy (TEM).

### *CPS conjugation to Tandem Core*™ *VLPs/Crm197*

CPS was oxidised and conjugated to carrier proteins by reductive amination as previously described [10]. Briefly, purified CPS was dissolved in 1 x PBS buffer at 5 mg/mL concentration and sodium periodate (NaIO_4_) was added to give a final 28 mM concentration. The reaction mixture was vortexed until dissolution of NaIO_4_ and then gently shaken for 3 h at room temperature. To remove the excess NaIO_4_, the reaction mixture was dialysed against MilliQ water in a dialysis tube with a molecular weight cut-off of 6-8 kDa and lyophilised. Oxidised CPS and the chosen carrier protein were dissolved in 1 x PBS buffer to give a final concentration ranging from 0.2 to 6 mg/mL. Then, 10 µL of 1 M NaCNBH_3_ solution in 10 mM NaOH was added for each mL of the reaction mixture, which was gently shaken at room temperature for 10 days. Afterwards, the reaction mixture was quenched by adding 10 µL of 1 M NaBH_4_ solution in 10 mM NaOH for each mL of the reaction mixture with shaking at room temperature for 3 h. The reaction mixture was dialysed against Milli-Q water in a dialysis tube with a molecular weight cut-off of 6-8 kDa and lyophilised.

### Analysis of conjugate vaccines - SDS PAGE and agarose gel analysis

SDS PAGE: loading buffer (10 μL) [Laemmli sample buffer (BIO-RAD): 25 % (v/v) glycerol; 62.5 mM Tris/HCl, pH 6.8; 2% (w/v) SDS; 5% (v/v) *β*-mercaptoethanol; 0.01 % (w/v) bromophenol] was added to protein (10 μL) in Milli-Q water. After heating at 100°C for 5 min, the samples were loaded onto a RunBlue precast gel (Expedeon) and run in RunBlue running buffer (Expedeon) [40 mM Tricine; 60 mM Tris/HCl; 0.1 % (w/v) SDS; 2.5 mM sodium bisulfite; pH 8.2] at 180 V for 53 minutes. Gels were removed from the case and stained for protein with Instant Blue (Expedeon).

Agarose gel: 1.2 % (w/v) agarose solution in TBE buffer [100 mM Tris-HCl; 90 mM boric acid; 10 mM EDTA] was poured into gel mould and left to set at 4 °C. Samples (20 μL) were loaded in DNA loading buffer (5 μL; New England Biolabs) and gels were run at 60 V for 120 mins.

### Negative stain TEM

Samples were diluted to approximately 0.1 mg /mL in 20 mM Tris HCl pH 8.0. A droplet (10 – 20 µL) of each sample was placed on a strip of Parafilm. Glow discharged formvar – carbon coated copper grids, 300 - 400 mesh (Agar Scientific) were place carbon side down on each sample. After 2 - 5 minutes adsorption, the grids were transferred to a droplet of 20 mM Tris HCl pH 8.0. The grids were then blotted and washed with 1 % (w/v) uranyl acetate in dH_2_O prior to staining with a droplet of 1 % (w/v) uranyl acetate for 10 seconds followed by blotting and air drying for at least 20 minutes. The dried grids were viewed in a transmission electron microscope.

### Immunogold labelling for TEM

Samples were diluted and adsorbed to glow discharged, formvar – carbon nickel coated grids as described above. After adsorption and washing in 20 mM Tris HCl pH 8.0, the grids were placed on droplets of blocking buffer (0.5 % cold water fish skin gelatin, 0.025 % Tween-20 in 20 mM TBS pH 8.0) and incubated for 45 – 60 minutes. CPS Primary antibody (Dstl) was diluted 1 in 50 in antibody diluent (0.05 % cold water fish skin gelatin in TBS pH 8.0). After blocking, the grids were incubated on droplets of diluted primary antibody for 60 minutes followed by washing by inversion over 3 successive droplets of antibody diluent. The washed grids were then incubated for 60 minutes on droplets of secondary antibody (gold conjugated anti-mouse diluted 1:25 in antibody diluent). Labelled grids were washed over 5 successive droplets of antibody diluent followed by a wash with 20 mM Tris HCl pH 8.0 and then stained with uranyl acetate as described above. As negative controls, samples were incubated with antibody diluent only in place of primary antibody and then processed as described for the other grids.

### Analysis of conjugate vaccines – protein and carbohydrate determination

Quantification of total heptose was carried out by phenol-sulphuric acid assay [36]. Total protein quantification was carried out by Pierce™ BCA assay [37].

### Conjugate vaccines (Antigen amounts and polysaccharide: protein ratios)

Due to inefficiencies of the reductive amination reaction, the amounts of CPS, Crm197 and VLP varied between vaccines but within each study the vaccine was standardised to CPS dose. The initial study at 103 and 240 x MLD utilised a CPS concentration of 10 µg per dose which was reduced in the later study to 4 µg per dose in order to discriminate between the vaccines (Table 1A and B). The vaccines contained 15 % (w/v) Alum per dose.

**Table 1:** (A) Antigen concentration (µg) per dose of each conjugate vaccine. (B) Ratio of polysaccharide to protein per dose of each conjugate vaccine.

| Challenge dose (MLD) | Antigen concentration per dose (µg) |  |  |
| --- | --- | --- | --- |
|  | CPS | Crm197 | VLP |
| 103 (IP) | 10 | 2.2 | 3.4 |
| 240 (IP) | 10 | 2.2 | 3.4 |
| 489 (IP) | 4 | 1.9 | 0.51 |

| Challenge dose (MLD) | Ratio of polysaccharide to protein |  |
| --- | --- | --- |
|  | Crm197 conjugate | VLP conjugate |
| 103 | 4.6 : 1 | 2.9 : 1 |
| 240 | 4.6 : 1 | 2.9 : 1 |
| 489 | 2.1 : 1 | 7.8 : 1 |

### Animal challenge

Groups of BALB/c female mice between 6 and 8 weeks old (Charles River UK) were acclimatised for two weeks prior to experimental start and vaccinated *via* the intra-muscular (IM) route on Day 0. Groups of control mice were given adjuvant only.

Vaccine boosts were given on days 14 and 28 and the mice were challenged *via* the intra-peritoneal route (IP) with 0.1 mL of *B. pseudomallei* K96243 at 7.66 × 10^4^, 1.79 × 10^5^ or 3.64 × 10^5^ CFU per mouse (103, 240, and 489 x MLD respectively). We have previously calculated the MLD in the BALB/c mouse model to be 744 colony forming units (CFU) by the IP route [10]. The mice were observed twice daily for a period of 35 days after challenge for signs of disease and culled at pre-determined humane end-points. All mice were tail-bled 2 weeks post-vaccination. All animal work was carried out according to the Animal (Scientific Procedures) Act 1986 and following challenge, the mice were handled within a containment level 3 half-suit isolator.

### Antibody analysis of animal sera

ELISAs were performed on sera collected 14 days after the third vaccination. 96-well plates were coated with purified CPS at 10 µg/mL in PBS (Dulbecco’s PBS 1x, -CaCl_2_, -MgCl_2_) and incubated overnight at 4°C. Each well was washed three times with PBS supplemented with 0.05 % (v/v) Tween-20 (Sigma). The wells were then blocked with 2 % (w/v) skimmed milk powder (Sigma) in PBS and incubated at 37°C for 1 hour. Following three further washes with PBS-Tween, two-fold dilutions of the mouse serum samples in PBS supplemented with 2 % (w/v) skimmed milk powder were made across the plate. Also included into separate wells was serum from Adjuvant vaccinated mice as negative controls. The plate was incubated for a further 1 hour at 37°C and washed three times in PBS-Tween. A 1:2000 dilution of isotype specific goat anti-mouse horseradish peroxidase conjugate (Biorad) in PBS supplemented with 2 % (w/v) milk powder was added to each well and the plate incubated at 37°C for 1 hour. Following six washes in PBS-Tween, 100 µL of Tetramethylbenzidine (KPL) substrate was added to each well according to the manufacturer’s instructions, and incubated at room temperature for 20 minutes prior to measuring the absorbance at 620 _n m_. A reading above the mean negative control (adjuvant only sera) plus three standard deviations was considered positive and the titre was determined to be the reciprocal of the final positive dilution.

### Enumeration of bacterial loads

Mice surviving to day 35 post-challenge were humanely culled and the spleens, livers and lungs removed aseptically into 2 mL PBS. The organs were homogenised into 900 µL PBS using a sterile 40 µm disposable cell sieve and the barrel of a sterile syringe. A dilution series (10^−1^ to 10^−7^) was prepared in 24 well-tissue culture plates (900 µL PBS per well with the addition of 100 µL of sample) and 250 µL from each dilution (neat to 10^−6^) were plated onto LB agar. Plates were incubated for 48 h at 37°C and the number of bacterial CFU was determined.

### Statistical analysis

For each animal experiment, appropriate group sizes were determined by a power analysis (allowing for sufficient power to elucidate an approximately 4-fold increase in hazard rate) and survival data was analysed by pairwise Log-Rank (Mantel-Cox) test [38] using the software GraphPad Prism (version 6.02). ELISA data was transformed to the logarithm of 10 and first analysed for differences in variance by the Brown-Forsythe test. Due to differences in variance, the ELISA data was analysed by the Kruskal-Wallis test and Dunn’s multiple comparisons.

## Results

### VLP production

Our efforts focussed on expression of the GD3K6D3G TCVLPs in *Nicotiana benthamiana*, using the pEAQ-*HT* expression system developed in Lomonossoff group [39, 40]. The tandem core pEAQ-*HT*-t-HBcAg-EL expression plasmid [30] was subjected to restriction digestion and subsequent re-ligation with primers coding the requisite sequence. Transformation of *Agrobacterium tumefaciens* by electroporation with the resulting pEAQ-*HT*-t-HBcAg GD3K6D3G plasmid was followed by agro-infiltration of *Nicotiana benthamiana* leaves with bacterial suspensions. Six days post infiltration (dpi), the leaves were harvested. After an extensive clean-up, VLPs were purified by gel filtration chromatography (Sephacryl S500) (Figure 1a). An estimation of the expression levels, based on comparison with standards, was made and shown to be in the region of 0.4 mg of protein per gram of plant tissue (ca 0.7 mg per plant). The samples were subjected to transmission electron microscopy (TEM) analysis, which showed high quality VLPs, correctly assembled with a homogeneous size of approximately 30 nm diameter and the characteristic HBc particle shape with small spikes on the surface (Figure 1b).

**Figure 1:**
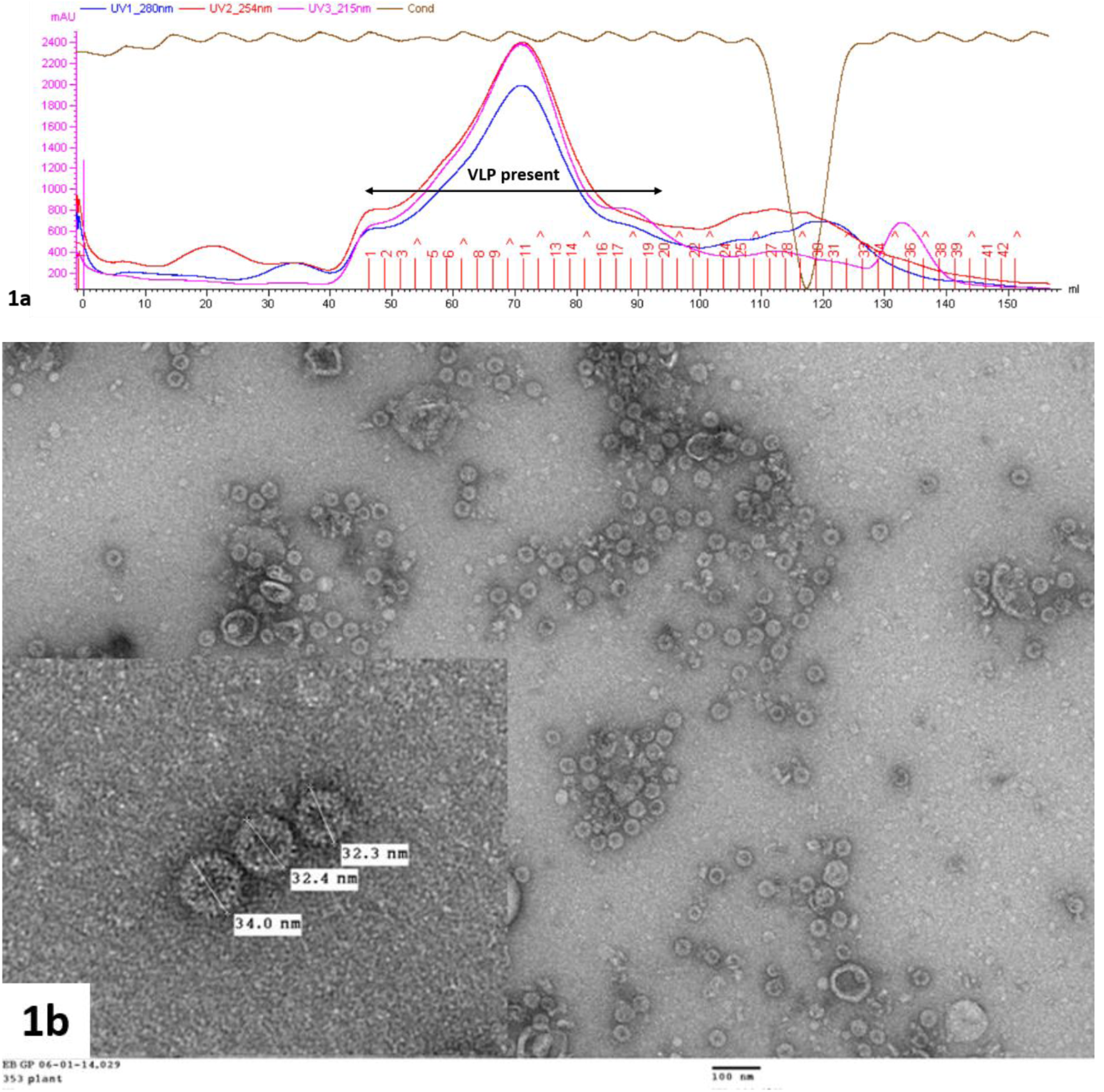
a) Chromatogram of Gel filtration elution for GD3K6D3G TCVLPs on Sephacryl S500 (VLP’s presence from fraction 1 to 20 was confirmed by TEM); b) TEM image of plant GD3K6D3G TCVLPs. Specimens were negatively stained with 2% (w/v) uranyl acetate; the scale bar in the large image indicates 100 nm.

### Glycoconjugate synthesis

To construct the glycoconjugates, extracted CPS from *B. thailandensis* strain E555 [32] was activated with sodium periodate and covalently linked to Crm197 or GD3K6D3G TCVLPs *via* reductive amination. Conjugation was confirmed by SDS PAGE and agarose gel with Coomassie staining (Figure 2A and 2B, respectively), which confirms the shift in molecular weight from unconjugated carrier protein to conjugate; and immunogold staining TEM (Figure 3A and 3B), which confirmed presence of CPS immunogenic epitope integrity through binding of an anti-CPS monoclonal.

**Figure 2:**
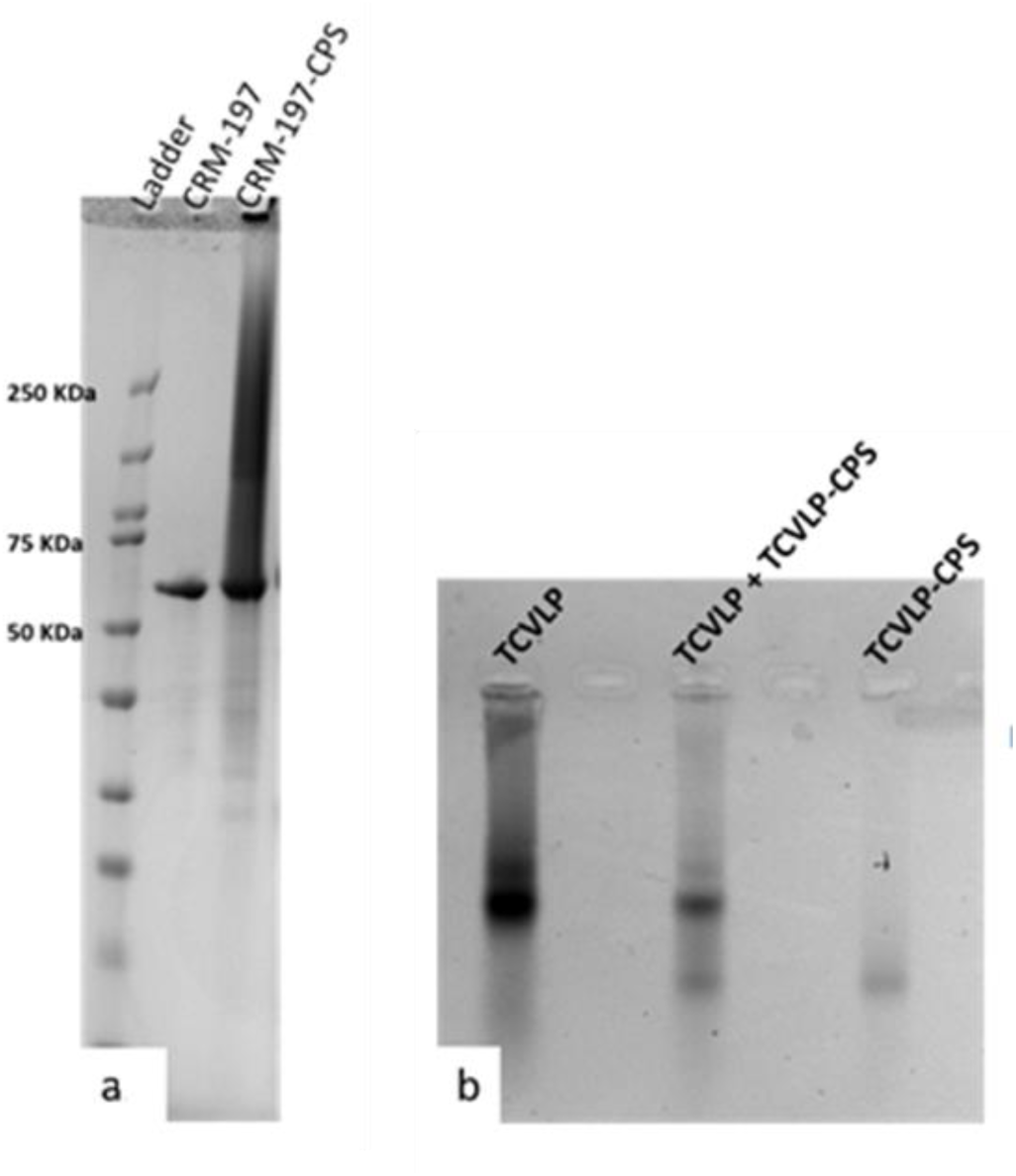
Gel electrophoresis of CPS conjugates. **(a) Crm197-CPS SDS PAGE, Coomassie blue staining;** L: ladder; lane 1: Crm197 (58.4 KDa); lane 2: Crm197-CPS. **(b): Plant TCVLP–CPS agarose gel (1.2% (w/v) in TBE), Coomassie blue staining.** Lane 1: Plant TCVLP; lane 2: TCVLP+ TCVLP-CPS; lane 3: TCVLP-CPS.

**Figure 3:**
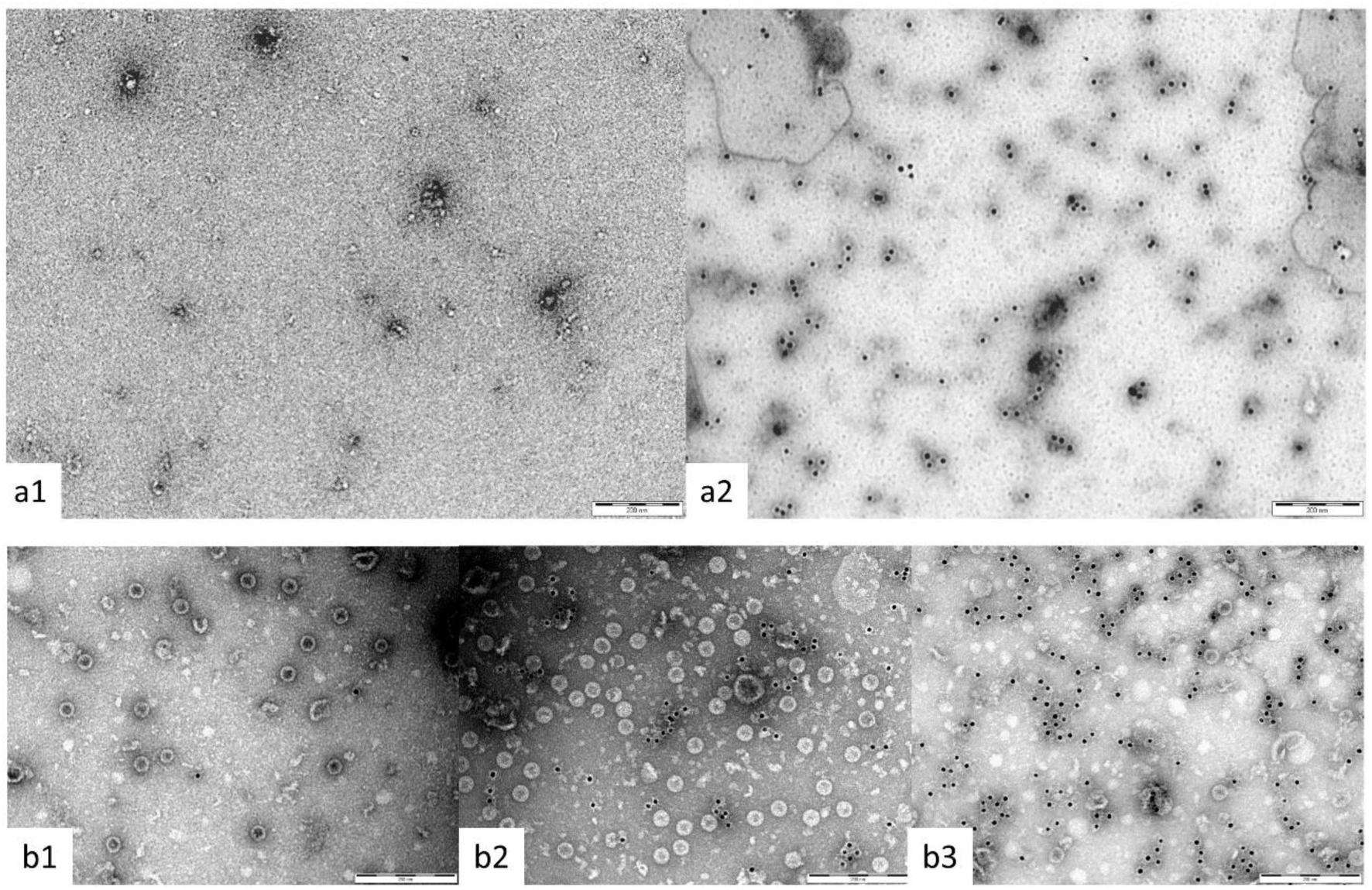
Immunogold staining TEM of CPS conjugates. A: Crm197-CPS immunogold staining TEM. (a1) Crm197 unconjugated control; (a2) anti-CPS mAb. **B: TCVLP-CPS immunogold staining TEM.** (b1) TCVLP unconjugated control; (b2) anti-VLP core mAb (10e11); (b3) anti-CPS mAb. Specimens were negatively stained with 2 % uranyl acetate; the scale bar indicates 200 nm.

### Comparison of glycoconjugate efficacy

The initial challenge study was designed to estimate vaccine efficacy and utilised two challenge doses of 7.66 × 10^4^ CFU per mouse (103 x MLD) and 1.79 × 10^5^ CFU per mouse (240 x MLD) of *B. pseudomallei* K96243 selected on the basis of previous work with CPS conjugates [10]. At 103 x MLD *B. pseudomallei* challenge there was no significant difference in protection between TCVLP-CPS vaccinated mice and those that received Crm197-CPS (Figure 4, A: p = 0.1385). Both conjugate vaccines gave significantly greater protection than CPS alone (TCVLP-CPS: p = 0.0005 and Crm197-CPS: p = 0.0117) but the majority of surviving mice were not clear of infection (Figure 4, D). The survival of mice vaccinated with CPS was not significantly greater than mice that received adjuvant alone (Figure 4 A: p = 0.6261). With a 240 x MLD *B. pseudomallei* challenge there was no significant difference in protective efficacy between the TCVLP-CPS and Crm197-CPS vaccines (p = 0.0982) but no survivors were clear of infection (Figure 4, B and D). In this instance, survival of mice vaccinated with CPS was not significantly different to mice that received TCVLP-CPS or Crm197-CPS (p = 0.0763 and p = 0.9394 respectively). In order to discriminate between the conjugate vaccines, the challenge dose was increased to 3.64 × 10^5^ CFU per mouse in the next study (489 x MLD). At this challenge dose, efficacy of the Crm197-CPS vaccine was significantly greater than TCVLP-CPS (Figure 4, C: p = 0.0185). In this instance, conjugate vaccine efficacy was significantly greater than CPS alone (TCVLP-CPS: p = 0.0049, Crm197-CPS: p < 0.0001) but no survivors were clear of infection (Figure 4, C and D). The survival data of the conjugate vaccines, CPS and adjuvant (Alum) from each challenge dose was collated into single survival curves for each antigen (figure not shown). There was no significant difference in efficacy between TCVLP-CPS immunised mice and Crm197-CPS immunised mice (p = 0.5458). Each conjugate was also significantly more efficacious than CPS alone (TCVLP-CPS: p < 0.0001, Crm197-CPS: p < 0.0001). In all mice that survived to study end, one mouse was clear of bacterial burden in the liver, lung and spleen and had received the Crm197 conjugate.

**Figure 4:**
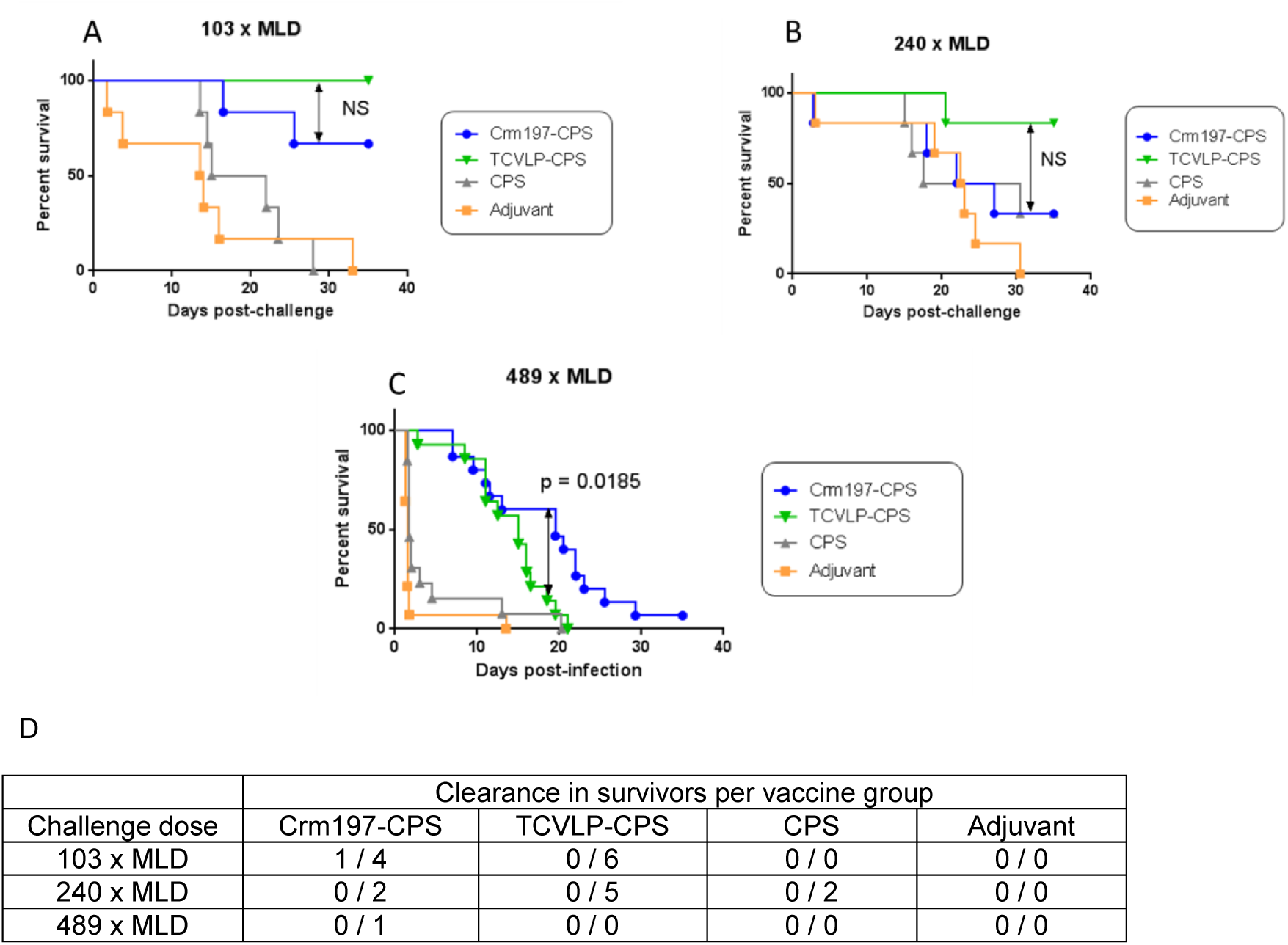
Efficacy comparison of vaccine antigens to 103, 240 and 489 x MLD *B. pseudomallei* K96243 challenge. Mice were immunised with CPS, Crm197-CPS, TCVLP-CPS, all formulated with Alum, or adjuvant (Alum), *via* the i.m. route on days 0, 14 and 28. Five weeks after the final immunisation, mice were challenged i.p. with 103 (A), 240 (B) or 489 (C) x MLD of *B. pseudomallei* strain K96243. Significance was determined by the log-rank (Mantel-Cox) test. 103 and 240 x MLD challenge: n = 6 mice per group. 489 x MLD challenge: n = 15 mice per group. (D) Bacterial clearance from the liver, lung and spleen in survivors per vaccine group between challenge doses. Individual tissues were mashed through a 0.45µm sieve filter and the resultant filtrate plated for bacterial counts.

### Comparison of glycoconjugate immunogenicity

ELISA analysis of serum obtained from tail bleeds after the third vaccination in each challenge study showed a CPS-specific IgM response from CPS-vaccinated mice and isotype switching to IgG in mice that received the conjugate vaccines. In all three challenge studies, mice vaccinated with Crm197-CPS had significantly greater CPS-specific IgM and IgG titres compared to mice vaccinated with TCVLP-CPS (Figure 5: 103 and 240 x MLD: IgM p ≤ 0.001, IgG p ≤0.0001. 489 x MLD: IgM and IgG p ≤ 0.05). For information, the individually-reported CPS-specific IgG and IgM responses generated in mice that received 10µg of CPS per dose were averaged by cage (n=6) for comparison to the cage-mean reported values for mice that received 4 µg/dose (Figure 5, black circles on 489 x MLD graph). Interestingly, the IgG responses from the Crm197 vaccinated mice were similar despite the difference in CPS amount, which may be due to the fact that the protein-polysaccharide ratio was similar between the two vaccines. In the TCVLP-CPS vaccinated groups, IgG responses were lower in mice that received the higher CPS concentration but this may be due to a lower polysaccharide-protein ratio in that vaccine.

**Figure 5:**
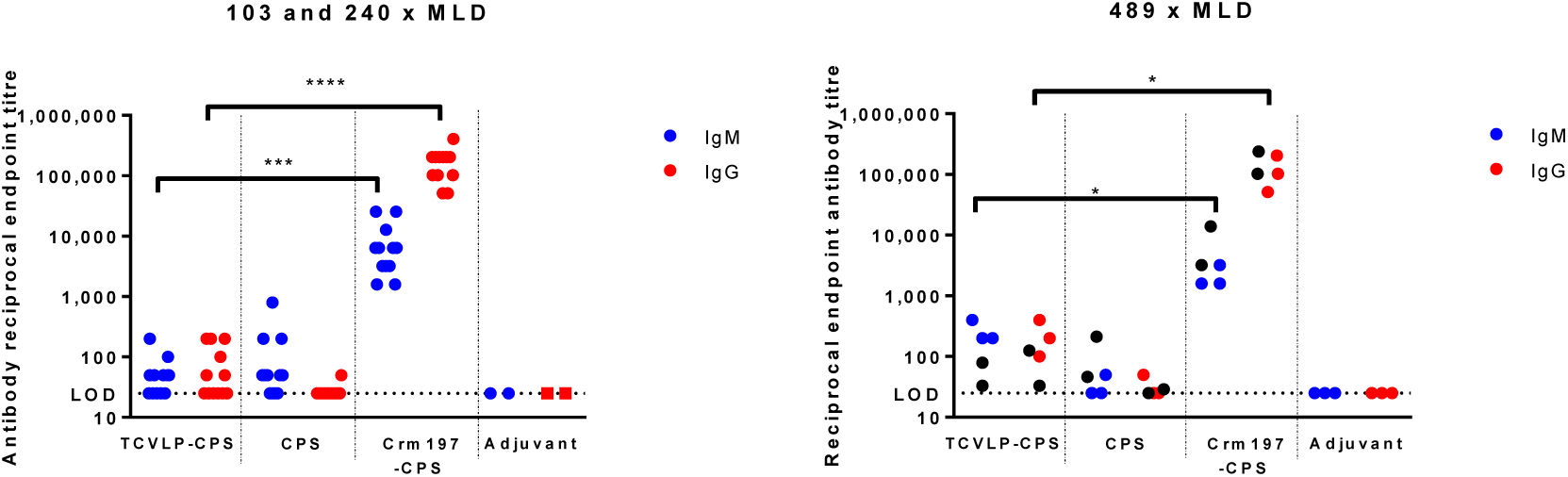
ELISA analysis of the CPS specific IgG and IgM antibody response from mouse sera obtained from the 103, 240 and 489 x MLD challenge studies. Mice were immunised on days 0, 14 and 28 *via* the i.m. route. Sera were obtained from mice 14 days after the final boost, and titres of IgG and IgM specific for CPS were determined by ELISA. 103 and 240 x MLD: individual symbols represent a single immunised mouse with exception of the adjuvant controls (n=6). 489 x MLD: individual symbols represent a cage of 5 mice. The data from the 103 and 240 x MLD graph is shown on the 489 x MLD graph for information (black circles) to demonstrate the similarity in CPS titres generated in mice that received 10 µg or 4µg CPS. Statistical significance was determined by Kruskal-Wallis test and Dunn’s multiple comparisons *p ≤ 0.05, ***p ≤ 0.001, ****p ≤ 0.0001.

## Discussion

Glycoconjugate vaccines have been instrumental in reducing disease incidence from several encapsulated bacteria including *N. meningitidis, S. pneumoniae* and *H. influenzae* type b, which use Crm197 as the carrier protein [16]. Despite the recent success of a Crm197 glycoconjugate protecting mice against *B. pseudomallei* challenge (14), there are concerns that exposure to multiple vaccines with the same carrier protein will result in immune interference leading to a possible reduction in vaccine efficacy (9, 15, 16, 17, 18). The aim of this work was to explore an alternative vaccine platform to Crm197 for conjugation to CPS with assessment of immunogenicity and protective efficacy in a murine model of melioidosis. Whilst Crm197 is an established carrier protein, VLPs have primarily been used as antigens to vaccinate against the virus they are derived from [41]. VLPs have been used experimentally as carrier proteins for antigens of viral or parasitic pathogens but their use for the treatment of bacterial infections is not well established [19, 20, 42]. Tandem Core™ is a genetically modified hepatitis B core protein which enables insertion of constructs into the major immunodominant region (MIR) of the core whilst remaining assembly competent [30]. In this study, genetic insertion of a construct containing six lysine residues flanked on either side by three aspartic acid residues into the MIR of core 2 allows for conjugation to *Burkholderia* CPS. The TCVLPs were produced from plants as plants have all the eukaryotic machinery for the correct post-translational modification and folding of the TCVLPs. Production in plants compared to other platforms (baculovirus, yeast and *E. coli*) is also advantageous in terms of cost, scalability and the low-risk of introducing human-relevant infectious agents, together with a high yield of production thanks to the transient expression technique [43]. To the author’s knowledge, this is the first documented use of a virus-like particle in melioidosis vaccine development.

The first animal study was performed in order to estimate vaccine efficacy. The protective efficacy of both TCVLP-CPS, and Crm197-CPS were assessed against two *B. pseudomallei* challenge doses of 7.66 × 10^4^ and 1.79 × 10^5^ CFU per mouse (103 and 240 x MLD respectively). In previous work, the MLD of *B. pseudomallei* K96243 infection in BALB/c mice by the intraperitoneal route was calculated to be 744 CFU [10]. At both challenge doses the absolute level of survival was greatest in mice that received the TCVLP-CPS vaccine although statistical significance over Crm197-CPS was not achieved. In order to discriminate between the vaccines, the challenge dose on the following study was increased to 3.64 × 10^5^ CFU per mouse (489 x MLD) and vaccine CPS content reduced from 10 to 4 µg/mouse. Under these conditions, efficacy of the Crm197 conjugate was significantly better than TCVLP-CPS. The difference in observed efficacy between these two conjugates could be due to potential differences in carrier immunogenicity or presentation of CPS to the immune system. Alternatively, as the vaccine doses were standardised on the basis of polysaccharide content, the amount of carrier protein was different between the two conjugates. This resulted in a different protein:polysaccharide ratio which is reported to affect vaccine immunogenicity in other conjugate vaccines (44).

The majority of mice surviving up to day 35 on all studies, with all vaccines, displayed continued bodyweight loss and clinical signs. At study end it is possible that these mice had entered the chronic phase of melioidosis and would have eventually succumbed to infection. It could be argued that these vaccine candidates had essentially extended the mean time to death as sterilising immunity was not achieved in all animals. While this may be true, the high challenge doses used across these studies were chosen to discriminate the protective efficacy between vaccine candidates only and are considered not realistic human exposures in natural scenarios. The pathology and clinical signs associated with the IP route of infection in these studies correlate well with those reported by Welkos *et al*., [45]. The most common pathological finding included abscess formation in the spleen and splenomegaly.

ELISA analysis of mouse sera taken after the third vaccination showed the presence of CPS-specific IgM antibody titres in mice that received CPS, and CPS-specific IgM and IgG antibody titres in the groups that received the conjugate vaccines. This finding was expected as the carrier protein stimulates development of T-cell dependent immunogenicity against the polysaccharide, which includes antibody isotype switching from IgM to IgG [46]. The significant difference in CPS–specific antibody titres generated between these vaccines may be attributable to differences in immunogenicity between the carrier proteins, the particulate nature of TCVLPs and different presentation of CPS to the immune system, or by variations in precise CPS loading between the conjugates. The significantly lower CPS-specific IgG antibody titres generated by a TCVLP-CPS conjugate suggests that titre is not indicative of vaccine efficacy at these challenge doses. This is unexpected as for nearly all licensed vaccines, prevention of infection correlates with the induction of specific antibodies. For three of the main bacterial pathogens that cause disease (*H. influenzae* type b, pneumococci, and meningococci), the correlates are presence of opsonophagocytic or bactericidal antibodies [47]. Furthermore, humoral immunity has been reported as an important mechanism of protection against *B. pseudomallei* infection [48]. One possible explanation is that low titres of CPS-specific antibodies are protective at lower challenge levels, which is feasible as it has been reported that correlates of protection are often relative to the challenge dose [49]. Alternatively, the TCVLP-CPS conjugate may generate low levels of high-affinity antibody which are protective at lower challenge levels; that the presentation of CPS to the immune system is different to Crm197-CPS; or that primary efficacy is *via* a different, perhaps cellular mediated, mechanism. This cellular mechanism in combination with a low level of antibody response may be superior at low challenge doses but at high doses bacterial numbers may overwhelm the humoral immune response, or deny the time needed for generation of a cellular response. Further investigation of potential cell-mediated effects from both conjugate vaccines warrants investigation.

The difference in protective efficacy seen between the Crm197-CPS conjugates used here and the ones used by Burtnick et al. [14] could be a result of several experimental differences. Firstly, the different animal models utilised by each lab. BALB/c mice used in this study are used primarily as an acute model of human melioidosis on the basis of proinflammatory cytokine release which correlates with disease severity [50, 51, 52, 53]. In contrast, C57BL/6 mice release lower levels of proinflammatory cytokines and therefore are used in the study of chronic melioidosis [51, 54, 55]. The challenge doses used in this study were also significantly greater, although disease progression from infection by the intraperitoneal route is less severe than inhalational challenge. Lastly, the use of alhydrogel and CpG by Burtnick et al. as opposed to alhydrogel alone as the adjuvant which was used in these studies could be beneficial given that CpG motifs have been shown to improve humoral and cellular immune responses [56].

The results from this study show that CPS conjugates utilising either Tandem Core™ or Crm197 as the carrier protein are effective vaccines for immunisation against melioidosis. The difference in generated IgG antibody titres between the conjugates warrants investigation.

## Funding Statement

This research was funded by the US Defence Threat Reduction Agency (DTRA), grant CBBAA12-VAXBT2-1-0032. Studies at the JIC were supported by the UK BBSRC Institute Strategic Programme on Understanding and Exploiting Metabolism (MET) [BB/J004561/1] and the John Innes Foundation. The funders had no role in study design, data collection and analysis, decision to publish, or preparation of the manuscript. WMR is a NIHR Senior Investigator and is supported by the UCLH NIHR BRC.

## Competing financial interests

G.P.L. declares that he is a named inventor on granted patent WO 29087391 A1 which describes the transient expression system used in this manuscript.

M. W. and R. A. F. declare that they are named inventors on granted patent WO 2015124919 A1 which describes the development of vaccines based on hepatitis b core antigens.

