## Supplementary information for "Assessments of hepatitis B virus-like particles and Crm197 as carrier proteins in melioidosis glycoconjugate vaccines"

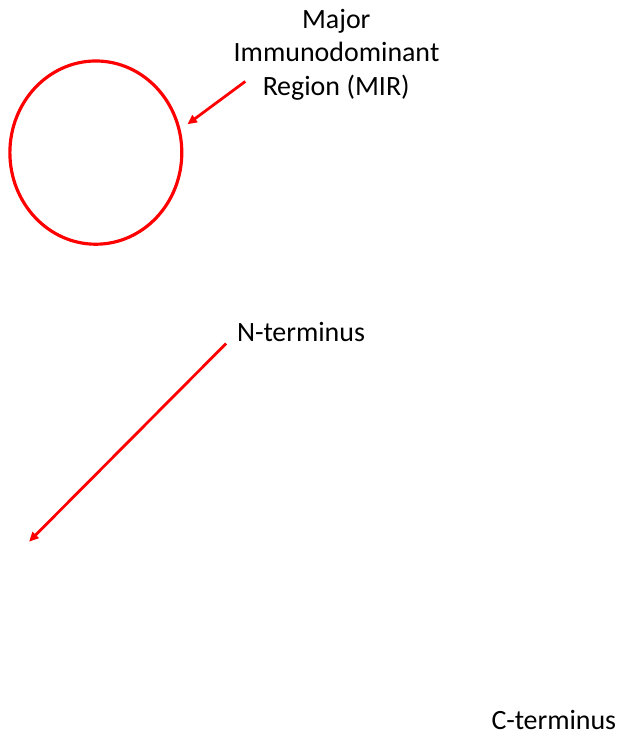

**Figure S1. Structure of HBcAg dimer (monomer in purple). Image reproduced from Wynne *et al. (***S. A. Wynne, R. A. Crowther and A. G. W. Leslie, *Mol. Cell*, 1999, **3**, 771-780.)

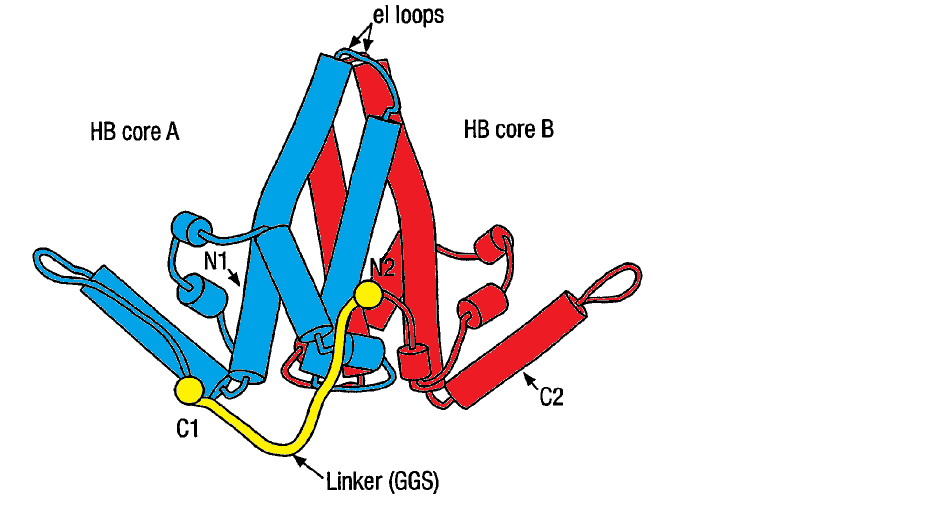

**Figure S2. Schematic of the tandem HBcAg core with two HBcAg monomers fused together with a flexible peptide linker highlighted in yellow (adapted from Gehin *et al* (**A. Gehin, R. Gilbert, D. Stuart, D. Rowlands, *US Pat.*, Hepatitis B core antigen fusion proteins, US 7270821, 2007)

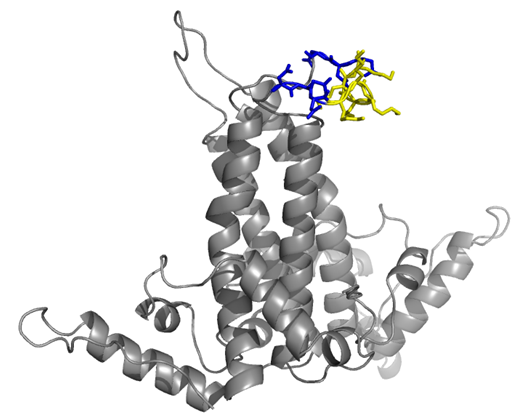

**Figure S3. Structure of wildtype HBcAg dimer with new protein insert (Lys - yellow, Asp - blue). Structure modelled using PyMol and PDB structure 1QGT.**
